# Performance and allocation can simultaneously shape behaviour in fire-disturbed populations of root vole

**DOI:** 10.1101/2023.08.04.551962

**Authors:** Jan S. Boratyński, Karolina Iwińska, Martyna Wirowska, Zbigniew Borowski, Paweł Solecki, Mariusz Ciesielski, Zbyszek Boratyński

## Abstract

Metabolic physiology and animal personality are often considered linked to each other, shaping ecological and evolutionary strategies along a life-history continuum. The energy allocation model predicts a negative while the performance model predicts a positive correlation between the rate of metabolic processes and behaviours, such as activity level. The models might operate simultaneously but depending on the context one can predominate over the other, determining expression of alternative pro- and reactive behavioural strategies. Large-scale fires, such as the one that burnt wetlands of Biebrza National Park (NE Poland), degrade natural habitats, affect amount of food and shelters and modify predatory-prey interactions. Fires pose also direct threat to survival of local populations, such as the wetland specialist root vole (*Microtus oeconomus*). We hypothesized that fire disturbance, by changing environmental context and selective regimes, determines mechanisms linking physiology and behaviour. Positive relation found among most studies, predicted by the performance model, would revert to negative relation, predicted by the allocation model, affecting animals ecological strategy in disturbed habitat. We repeatedly measured maintenance and exercise metabolic rates and activity behaviour on voles from post-fire and unburnt populations. Repeatable maintenance metabolism and activity level were positively correlated, but more labile exercise metabolism did not explain behaviour. The correlations were not strongly affected by fire disturbance, but voles from post-fire habitat had higher maintenance but not maximum metabolism and moved shorter distances than individuals from unburnt area. The results suggest that performance model predominates, while habitat disturbance might reveal some allocation constraints on physiology-personality linkage.

**Summary statement:** Contrasting ’allocation’ and ’performance’ models, for energetics-behaviour linkage, were tested in context of fire-disturbance. Positive (performance) correlation predominated but animals from burned habitat had elevated metabolism and suppressed exploration (allocation).

## Introduction

Animal behaviour is a first-line response to selective pressure and as such has strong ecological and evolutionary consequences (Wolf and Weissing 2012).

Particularly the animal personality, understood as temporally persistent among individual differences in behaviour, is considered an important evolutionary trait (Réale et al., 2007). The variation of animal personalities is considered as a continuum on which individuals can be assigned as more proactive or more reactive (Koolhaas et al., 1999; Sih et al., 2004ab; see examples in: Šíchová et al., 2014). The reactive animals are unexplorative, shy and less likely to take risks, while proactive ones are explorative, bold and aggressive. Animal personality traits can be heritable (Dochtermann et al., 2015), may affect individual fitness (Smith and Blumstein 2008; Moiron et al., 2020 see also: Haave-Audet et al., 2022) and can also undergo correlative evolution along with morphological and physiological traits (Gębczyński and Konarzewski 2009a; Careau et al., 2011; Kern et al., 2016; Maiti et al. 2019; Zablocki-Thomas et al., 2019).

Basal metabolic rate (BMR) - the lowest metabolism at rest in endothermic animal (generating heat internally) under homeothermy (high and stable body temperature) - was for two decades studied in context of animal personality (Careau et al. 2008; Biro and Stamps 2010). Among others, this is because BMR is considered a major component of the animal daily energy expenditure of homeothermic animals (Ricklefs et al., 1996; Speakman, 1999; Portugal et al., 2016; Abdeen et al., 2021).

Since animal energy budgets are finite, the ‘compensation’ model predicts that the individual self-maintenance costs must limit other functions (Olson et al., 1992; Nilsson 2002; Blackmer et al., 2005) including expression of costly behaviours (Careau et al. 2008; Careau and Garland 2012). However intuitive, this model is rather rarely supported across behavioural studies (review in: Mathot et al., 2019, see also: Krams et al., 2017; Careau et al., 2019; Biro et al., 2020; Peña-Villalobos et al. 2020). The variation in BMR correlate with the variation in the mass/size of organs involved in the transformation of food into usable energy (Meerlo et al., 1997; Chappell et al., 1999; Russell and Chappell 2007: Raichlen et al., 2010). Thus, the alternative ‘performance’ model predicts that proper metabolic machinery and related high metabolism is required to support energetically costly behaviours (Careau et al. 2008; Biro and Stamps 2010). BMR is also considered a by-product cost of the evolution of endothermy - it increased as a correlative response to selection for maximum metabolic rate (MMR) or aerobic capacity during exercise (Bennett and Ruben 1979; Nespolo et al., 2017 but see also: Wone et al., 2015; Enriquez-Urzelai and Boratyński 2022). Surprisingly, only a few behavioural studies explored the relationship between individual variation in behaviour and maximum aerobic capacity during exercise (Jónás et al., 2010; Killen et al., 2014; Rupia et al., 2016; Behrens et al., 2020, review in: Biro et al., 2018; see also: Boratyński et al., 2020). Both resting and active metabolism are considered to be repeatable, heritable, affect animal performance and have fitness consequences (Dohm et al., 2002; Nespolo and Franco 2007; Boratyński and Koteja 2010; Auer et al., 2016; Boratyński et al., 2018; Pettersen et al., 2018; Arnold et al., 2021). Its variation at the population level might reflect the selection process (Boratyński and Koteja 2009; Wone et al., 2015; Sadowska et al., 2015) but it can also be altered as a result of developmental plasticity (Norin et al., 2019) or reversible flexibility (Swanson et al., 2017) caused by environmental perturbations or disturbance.

Anthropogenic habitat destruction is an important global cause of biodiversity loss, as it also degrades local biota resilience to climate change. Among others, due to global changes and the associated drying climate, large-area man-made fires become more frequent even in habitats for a long time considered fire-free (Bowman et al., 2020). In the fire-prone ecosystems animals evolved perception and proper response to smoke, sound and view of fire, allowing them to survive such disaster (Álvarez-Ruiz et al., 2021; Batista et al., 2023). However, many species are not adapted to fire which can be a serious threat to such fire-naive populations (Nimmo et al., 2021, 2022; Batista et al., 2023). Small terrestrial mammals seem most vulnerable to fire due to their limited movement ability (Erwin & Stasiak 1979; Tomas et al., 2021). Such populations may even temporarily go extinct at large areas under very severe fires – their recovery is possible only through recolonization. Recent macroanalysis suggests that mortality during fire is related to its severity, and that the proportion of killed individuals is usually not very high (Jolly et al., 2022). However, this study contains the existing data from fire-prone ecosystems occupied by well fire-adapted animals, or impacts of low-severity fires. The authors main conclusion was that our knowledge on this topic is still scarce, and they highlight the critical need to study the high severity fire in fire-naive animal populations (Jolly et al., 2022).

Post-fire environment can be considered challenging for the animals. Several factors such as reduced amount of food, natural vegetation cover and shelters, as well as increased predation may result in strong selective pressure on animal behaviour and physiology (Sutherland and Dickman 1999; Doherty et al., 2022; Jolly et al., 2022).

Despite the limited studies of animal behaviour (but not necessarily animal personality) in the context of fire-related disturbance, some general predictions can be adapted from predator-prey interaction theory (Doherty et al., 2022; Michel et al., 2022). The successful behavioural response to fire relies on animal vigilance, cue perception, assessment and proper response similar like in case of predator-prey interaction which is likely also affected by the evolutionary fire history of the population (Michel et al., 2022). In post-fire environment, the prediction of prey behavioural response can however depend also on the behavioural reaction of the predators (Doherty et al., 2022). Those frameworks give plausible scenarios, however, they seems to ignore the association predicted by ‘allocation’ and ‘performance’ models association between animal personality and physiology that could also modify behavioural responses in post-fire environments. Likewise, only a few studies on animal physiological adaptations were conducted in fire-ecology (review in: Geiser et al., 2018), and general physiological implications of dealing with post-fire disturbed habitats are still unsettled. Of help could be well-formulated models linking behaviour and physiology rooted in evolutionary ecology (see in: Careau and Garland 2012). Since the fire is strongly modifying the environment (e.g. resources and thermal conditions), and the animal physiology is environment dependent, then the correlated behavioural changes at post-fire habitats would not only result from predator-prey interaction.

Large fire happened at the open landscape of wetland-grassland in Biebrza Valley (Biebrza National Park; Poland) in the spring of 2020. In a single week a one fourth of the habitat and, along with it, a significant part of the population of root vole (*Microtus oeconomus*) was fully demolished – grass covers were burnt and voles nests destroyed. Initial screening conducted one month after the fire indicated a lack of any rodents in the burnt zone. This created an unique opportunity to test whether the animal personality, morphological and physiological traits respond to challenges that fire-naive animals meet at post-fire habitats. We hypothesized that during recolonization of burnt habitats there is a high selective pressure on animal behaviour (an effect of increased predation) and on physiology (an effect of reduced resources). We predicted that individuals successfully colonizing post-fire habitat will be selected for (repeatable) reactive behaviour (shy/less explorative), when compared to voles from unburnt habitats. Since animal personality traits are expected to covary with physiology we also predicted its correlated differences between burnt and unburnt sites. Specifically, we predicted that in burnt habitats, due to lowered resources availability (food and shelters), voles personality traits will be shaped by their physiology according to the ‘allocation’ model. In contrast, we expected that ‘performance’ model could operate in voles from unburnt habitats. Thus, we expect to find a positive correlation between proactive behaviour (exploration/boldness) and BMR in animals captured at unburnt, and the opposite relationship in voles from burnt habitats. Since the upper limit to aerobic metabolism may be a better measure for individual performance, we expected an even stronger positive correlation between proactiveness and maximum metabolic rate in voles from unburnt habitat. Although, we expected negative correlation between BMR and proactiveness in voles from burnt habitat, we did not exclude the possibility that a positive correlation persist between activity behaviour and maximum metabolism. To verify these predictions, we tracked the voles recolonization at the burnt sites in reference to unburnt plots in vicinity from late spring till autumn. After population restored at the burnt part in autumn (6-7 months after the fire), we quantified behavioural traits and measured metabolic rates on voles from both burnt and unburnt habitats. To verify if phenotypes reflect consistent individual differences, we repeatedly captured and measured marked animals until spring.

## Material and methods

### Study site and animals

The Biebrza Valley is a large complex (136 900 ha) of sedge wetlands in NE Poland (Żurek 2005) that can be divided into three connected parts Upper, Middle and Lower Basins. The whole valley is a mosaic landscape, however, its core zone is temporally flooded and dominated by homogeneous water resistant low vegetation of tussock sedge *Carex* spp. (Borowski et al., 2021). Root vole is a dominant rodent species in this habitat forming ∼90% of small mammals community (Borowski 2002). This species utilizes clumps of sedge as a nesting place and also depend on them as its main food source (Borowski et al., 2021).

Seasonally variable water level is usually the highest during spring (after winter snow melts) and floodings are frequent. Decline in groundwater level due to water management in the past together with current climate changes (as lower and unpredictable snow falls in winter) currently result in increased frequency of fires in this area. The biggest human-caused wildfire took place in central part of Biebrza Valley at an area of >5500 ha (Fig. 1). In result ∼90% of surface green-cover in Middle Basin burnt, as well as a habitat of root voles and their nests (Fig. 2).

**Figure 1.**
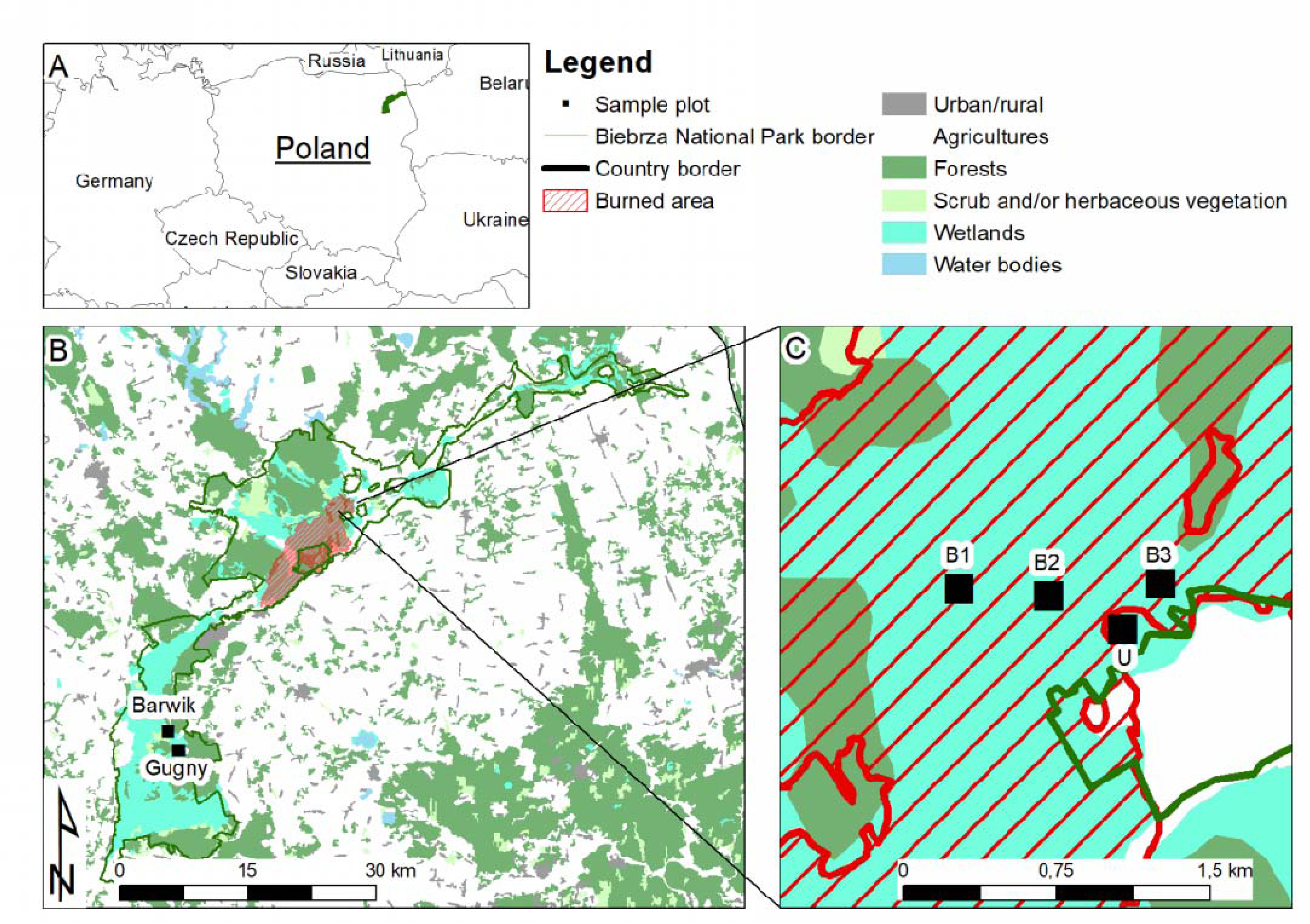
Map of the study area. Location of plots at burnt habitat (B1, B2, B3) in Middle Basin, control plot at close fire vicinity (U) and control plots at Lower Basin (Barwik and Gugny) are presented.

**Figure 2.**
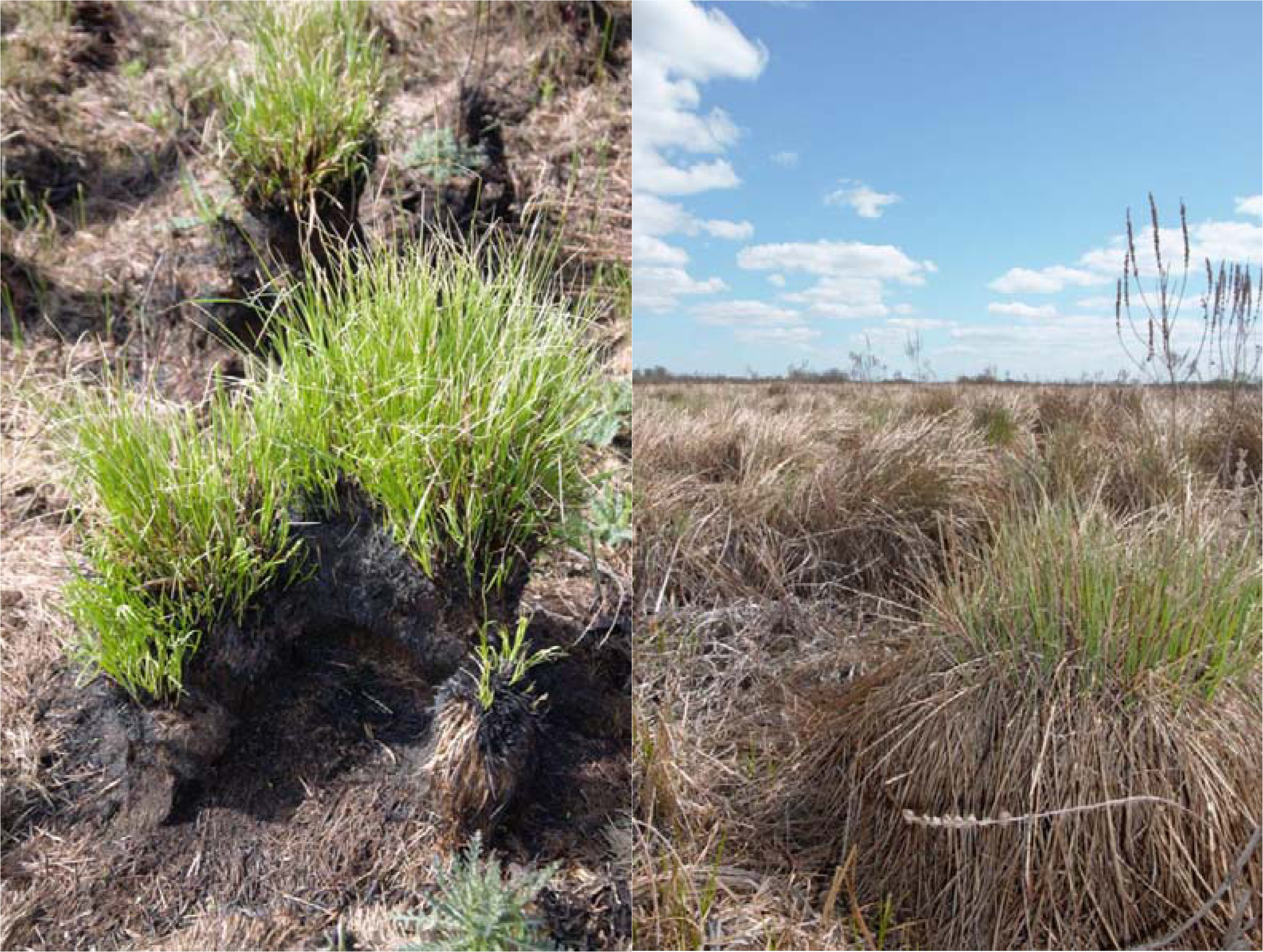
Examples of clumps of sedges that used as nests by root voles. Left photo represents burned root vole nest and surrounding one month after fire. Right photo show part of the undisturbed habitats with clumps of sedges and accumulated dry plant mass (photo by Jan S. Boratyński).

The study was conducted at the burnt and unburnt parts of the valley in Middle Basin (53°35’N, 22°53’E), and at the permanent research plots located in Lower Basin (53°21’N 22°33’E). Three plots were established at burnt habitat (B1, B2, B3) and a single one at unburnt habitat (U; Fig. 1) at Middle Basin. For the reason the unburnt part at Middle Basin was very limited or inaccessible (lack of roads at very swampy areas), we also captured voles at our permanent plots located at Lower Basin (∼30km apart; Fig. 1).

The field and laboratory experimental procedures were approved by the Second Local Committee for Ethics in Animal Research in Warsaw (decision no. WAW2/112/2020) and the Biebrzański National Park (decision no. 38/O/2020). Animals were captured using wooden live-traps forming 0.6 ha plots of 10 × 10 m grids (two traps placed at each point). At study plots located in Middle Basin animals were trapped monthly (five subsequent days) from the summer of 2020 to track the recolonization process. In October and November of 2020, animals captured at all six study plots were transported to the laboratory at the Field Station of the Faculty of Biology at University of Białystok located in Gugny village (53°21’N 22°35’E). In the laboratory voles were kept individually in standard rodent cages (model 1264, Tecniplast, Italy) lined with hay, equipped with paper tubes as shelter and with access to rodent dry food (Megan, Poland), carrots and water *ad libitum* at an ambient temperature of 15_ and natural photoperiod. For each individual behavioural, morphological and physiological traits were measured. After three days spent in the laboratory, when all procedures were finished, animals were released at the place of capture. Before release, the animals were individually marked using radio-frequency identification tags (RF-IDW-2, CBDZOE, Poland). We continued capturing and measuring voles from Middle Basin in late November 2020, late January and March- April 2021 to estimate the repeatability of all measured traits.

### Open field test

On the day following capture, in the evening (∼1h after sunset), voles were exposed to an open-field test (OFT) in order to quantify their behaviour. OFTs were conducted during nighttime to fit it into the period of the main activity of our model organism. Self-constructed (opaque polyvinyl chloride plastic) arena (1 × 1 × 1 m) placed in a dark room was illuminated by 5.5 W bulbs (470 lm; Leroy Merlin Poland, Warsaw) painted red and mounted outside above each corner (at 2 m height). As a result, central part of the arena was illuminated at a level of 50 lxs. Before each OFT the arena was cleaned with 70% alcohol and dried. At the beginning of OFT, animals were carefully placed in the corner using a one meter tall pipe which was removed immediately when the animal reached the floor. Recordings of animal behaviour were carried out automatically with digital camera (Hero 5, GoPro Inc., US) mounted at 2.0m above the central part of arena. The observers (KI and JSB) were leaving the room and tracking the recordings (which lasted five minutes) with a smartphone using GoPro App (GoPro Inc., US) to avoid disturbance.

### Respirometry

To measure metabolic rate, indirect calorimetry using open-flow respirometry was used. The air was pulled from outside, warmed up to 28-30 _ and dried (silica gel 2-5 mm, Chempur, Piekary Śląskie, Poland). Maximum metabolic rate (MMR; aerobic capacity) was measured on the following night (starting 1h after sunset) after OFTs. MMR was measured during forced swim at Ta = 28 _, slightly below lower critical temperature for thermoneutral zone to avoid overheating the animals. In this case we used single-chamber (∼1 L) one-line system with flow rate (∼1 L · min^-1^) set and measured incurrently using mass-flow meter (ERG-1000, Warsaw, Poland). The chamber (placed in water bath) was filled with water leaving only 250mL of empty space inside for swimming voles. In this system single oxygen analyzer (FC-10a, Sable System Int.) was used and air leaving the chamber was dried and set to pass it through analyzer at the rate of ∼100 mL · min^-1^. Basal metabolic rate (BMR, physiological maintenance costs) measurements were performed during the day following MMR measurements. To measure BMR in eight animals at once we used a two-line system with two oxygen analyzers (FC-10a, Sable System Int.). Four chambers (∼700 mL) were placed at Ta = 30 _ (water bath) connected to each line and air was subsampled at rate of 100 mL · min^-1^ excurrently with use of two-line computer-controlled multiplexer (MUX, Sable System Int., USA). This allowed to sample two animals simultaneously and switch between channels and baseline automatically. The flow rate (∼400 mL · min^-1^) of dried air leaving the chamber was measured at each line using two mass-flow meters (ERG-1000, Warsaw, Poland). All electronic elements of the systems were connected to PC via analog-digital interface (UI-2 SSI) and data were collected using ExpeData (SSI) software at 1 Hz. Each animal was sampled for 2 minutes every 11 minutes. The mass-flow meters were calibrated against soap bubble flowmeter (model: Optiflow 570, Humonic Instruments Inc., USA). Before each metabolic rate measurements *m*_b_ was obtained to nearest 0.1g (ScoutPro 200, Ohaus, Parsippany, NJ, USA).

### Data processing

Behavioural variables as the total time spent rearing or grooming were counted manually (single observer MW), the total distance moved and the time spent in different parts of the arena (center, corners or near walls) were measured automatically with a trained neural network using DeepLabCut software (Mathis et al. 2018). Based on obtained row data we calculated frequencies of time spent rearing, grooming and frequencies of time spent in the center or corners.

Oxygen consumption during BMR and MMR measurements were calculated using equations 11.2 and 10.2 after Lighton (2008), respectively. We assumed a respiratory exchange ratio equal to 0.8. Metabolic rate during MMR was measured as maximum 30 second average oxygen consumption during swimming. Metabolic rate during BMR measurements was calculated as the lowest 20 second readings of the most stable 40 second readings of the 1 minute from the end of each sample. BMR was assumed as the lowest reading obtained for the individuals during a measurement session (22 samples were obtained during each ∼4 h session).

### Statistics

All statistics were calculated in R program (ver. 4.3.1). In the first step we estimated repeatability (τ) for all measured traits using function ‘rptR’. In all models for estimating τ’s, individual *m*_b_ was included as covariate, sex as factor and animal ID as a random effect. For behavioural and physiological traits, we also ran models where subsequent test (henceforth: test sequence) was included as an additional factor. The Gaussian distribution was assumed when τ was estimated. However, because of the heavy right skewed distribution of model residuals for frequency of time spent rearing, frequency of grooming and frequency of time spent in center, those variables were square-root transformed prior to the analysis. We estimated τ’s based on data available for 29 animals measured at least twice (four individuals were measured three times). However, we also kept the individuals measured only once (101 individuals) to improve the general fit of the dependent variable to explanatory variables during the modeling procedure. Between subsequent measurements in this analysis, individual voles spent 18-157 days in the field conditions (mean ∼61 days).

Linear mixed effects modeling (LME) procedure was used to compare only the repeatable traits (see results) as *m*_b_, BMR, MMR and distance moved by voles captured at burnt and unburnt sites using the function ‘lme’. Only naive individuals (first time measured) captured in autumn (October and November) were included in this analysis. Dependent variables were not significantly correlated and thus we used separate univariate LMEs during analysis. In all LMEs, sex and fire-status of place were included as factors and site ID as a random effect. In the LME for BMR, MMR and distance moved *m*_b_ were included as a covariate. In the LME for MMR residual BMR (obtained from its linear relationship with *m*_b_) was included as a covariate. In the LME for distance moved, residual BMR and residual MMR (obtained from its linear relationship with *m*_b_) were included as covariates. Interactions between fire- status and residual BMR or residual MMR tested in the initial LME for distance moved were finally excluded from the model due to its insignificant effects (see results). Due to heteroscedasticity of model residuals, the distance moved was square root transformed prior to the analysis. All models were tested using analysis of variance (type II and III of sum of squares for main effect models and models with interactions were used, respectively). We calculated estimated marginal means using the function ‘emmeans’ to compare fixed effects terms.

## Results

In June, during trapping in Middle Basin, the density of voles was lower at three burnt sampling plots than at the control plot (Fig. 3). The density of voles at the control plot in Middle Basin was similar to those at Lower Basin plots (Fig. 3). In the following months population density tended to increase but it was always lower at burnt plots when compared to controls (Fig. 3).

**Figure 3.**
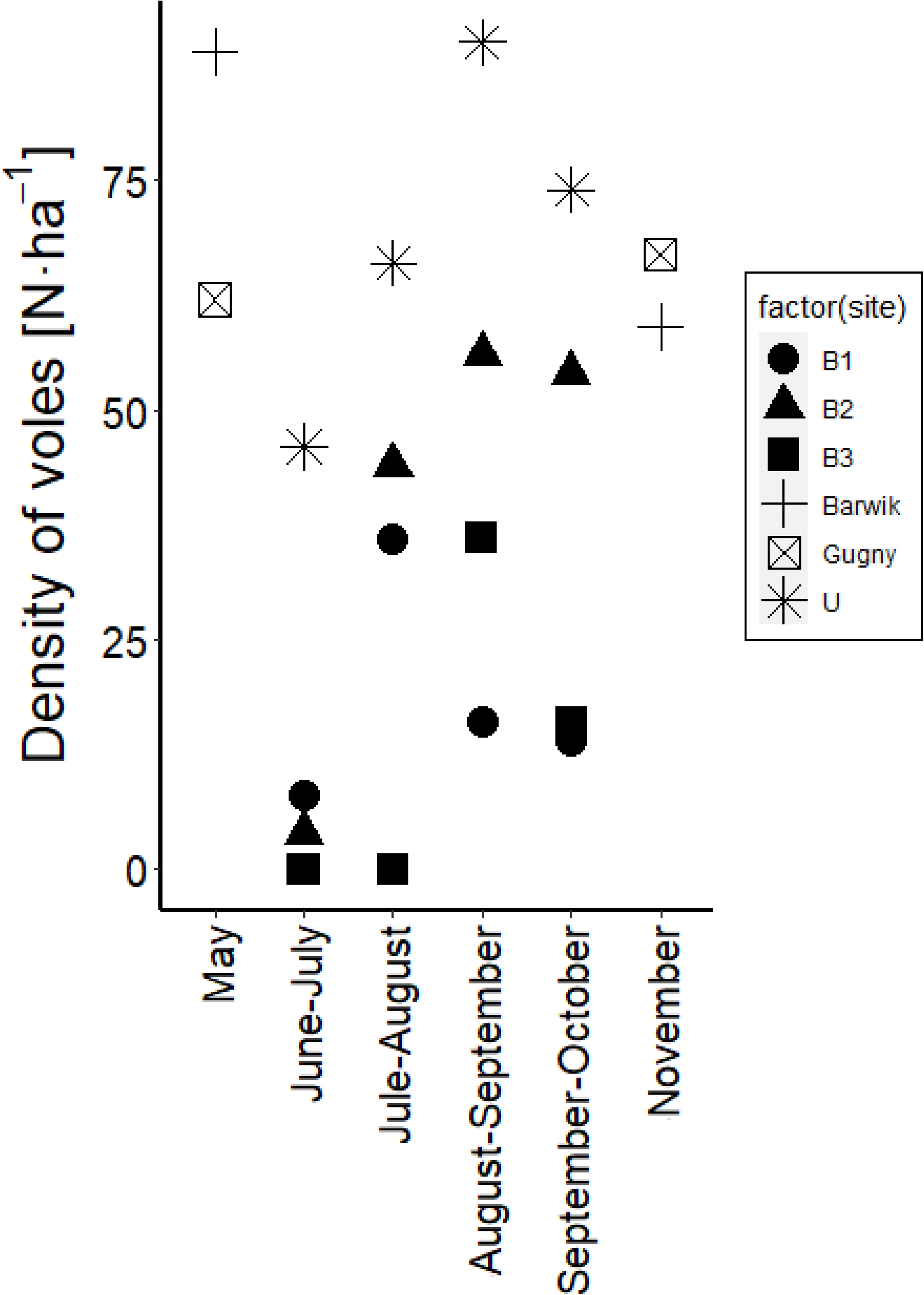
Changes of root vole population density. Dark symbols (circle, triange and square) indicate vole densities at burnt (B1, B2, B3) and the others (cross, star and crossed-sqare) indicate vole densities at unburnt sites in Middle (U) and Lower Basin (Barwik and Gugny). Population density was calculated as the minimum number of alive individuals per ha.

When sex was maintained as factor during analysis, *m*_b_ was found repeatable (Tab. 1). When adjusted for variation in *m*_b_ and sex, τ was found significantly higher from zero for BMR (Tab. 1). Sex and *m*_b_-adjusted MMR was found significantly different from zero only when test sequence was included in the analysis (Tab. 1).

**Table 1.**
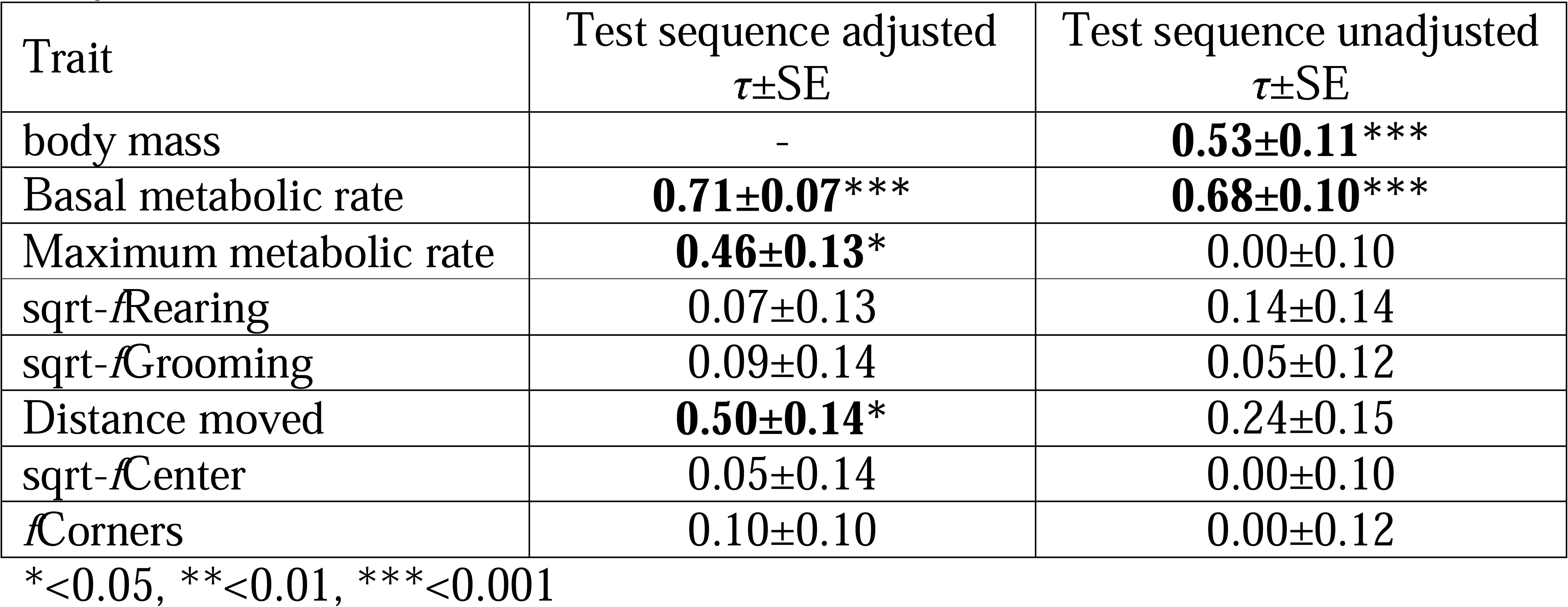
Repeatabilities (τ) estimated and their standard errors (SE) for behavioural, morphological and physiological traits in root voles. Estimates for traits adjusted for test sequence (when naive animals were tested for the first time or repeated) order of measurements was included (adjusted) and ignored (unadjusted). Basal and maximum metabolic rate and behaviours were adjusted for sex factor and body mass covariate (for details see material and methods).

The distance moved during OFT was a significantly repeatable trait in root voles when sex, *m*_b_ and test sequence were included into analysis (Tab. 1). Other behavioural categories included in analysis were found not to differ consistently among individuals.

There was no significant differences for *m*_b_ between sexes (Tab. 2). There was also no differences for *m*_b_ between place of capture (Tab. 2). BMR was positively correlated with *m*_b_ but did not differ between sexes (Tab. 2). Sex and *m*_b_-adjusted BMR was higher in voles captured at burnt plots when compared to those from unburnt locations (Tab. 2). There was no interaction for MMR between fire-status and residual BMR (F_1,114_ = 2.35, *P* = 0.128). Neither sex nor fire-status affected MMR, which correlated positively with *m*_b_ but not with residual BMR (Tab. 2). Sqaure-root transformed distance moved was negatively correlated with *m*_b_ and did not differ between male and female voles (Tab. 2). The interactions between residual BMR or residual MMR and fire-status were insignificant in a model for square-root transformed distance moved during open-field test (residual BMR: F_1,112_ = 1.73, P = 0.192, residual MMR: F_1,113_ = 2.95, P = 0.089). Distance moved (*m*_b_-adjusted, square- root transformed) correlated positively with residual BMR and was lower in voles captured at post-fire plots when compared to those from unburnt sites (Tab. 2; Fig 4b).

**Figure 4.**
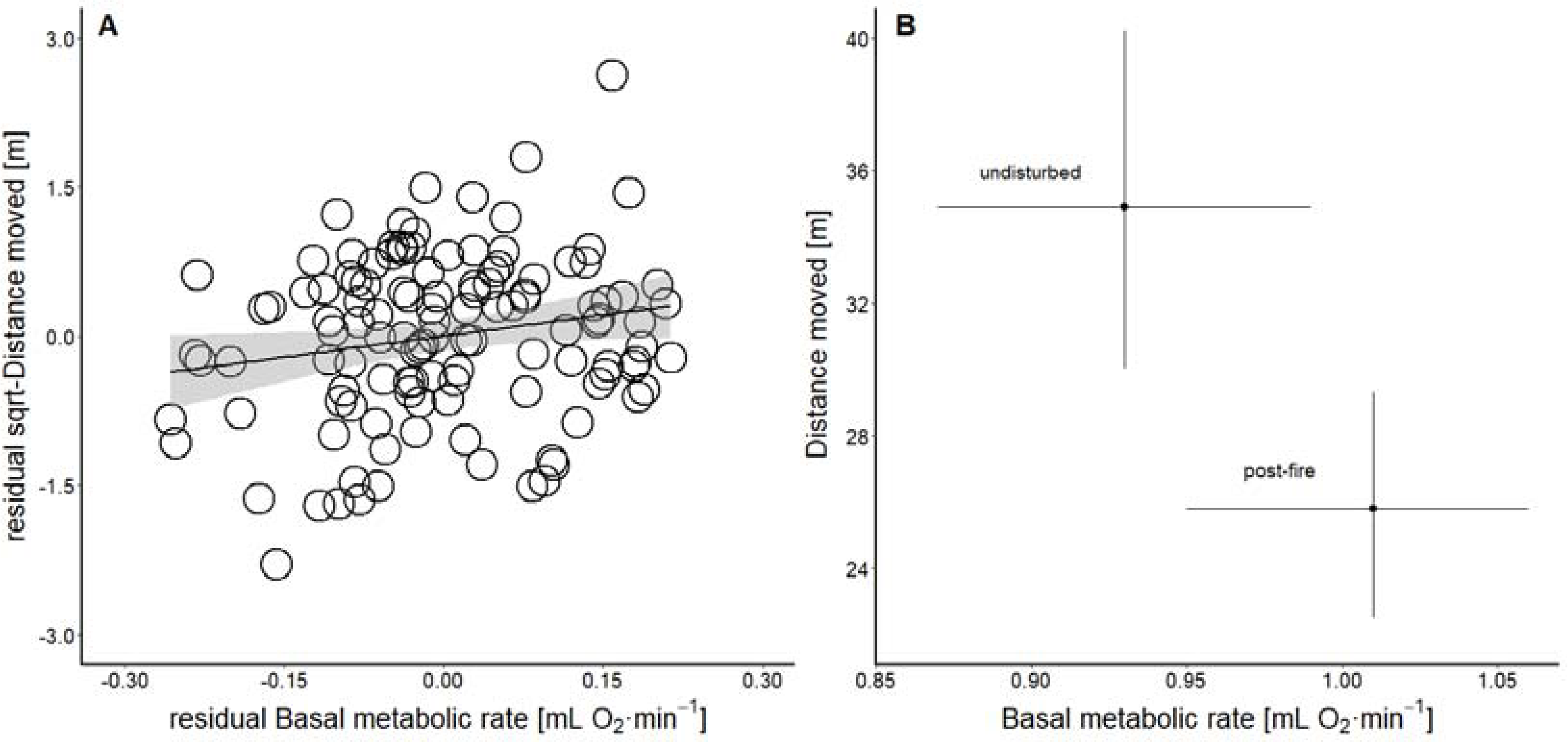
Correlation between basal metabolic rate and distance moved. A) Relationship between residual basal metabolic rate and residual square-root transformed distance moved (both corrected for body mass and fire-status) in root voles during open field test. B) Estimated marginal means for basal metabolic rate and distance moved during an open field test in voles captured at the post-fire zone and undisturbed habitat obtained from linear mixed effect models.

**Table 2.**
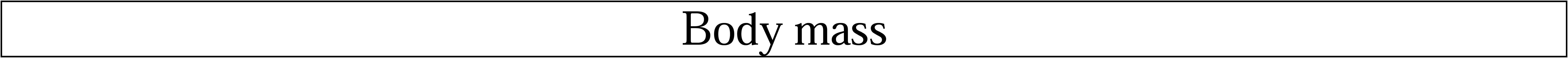
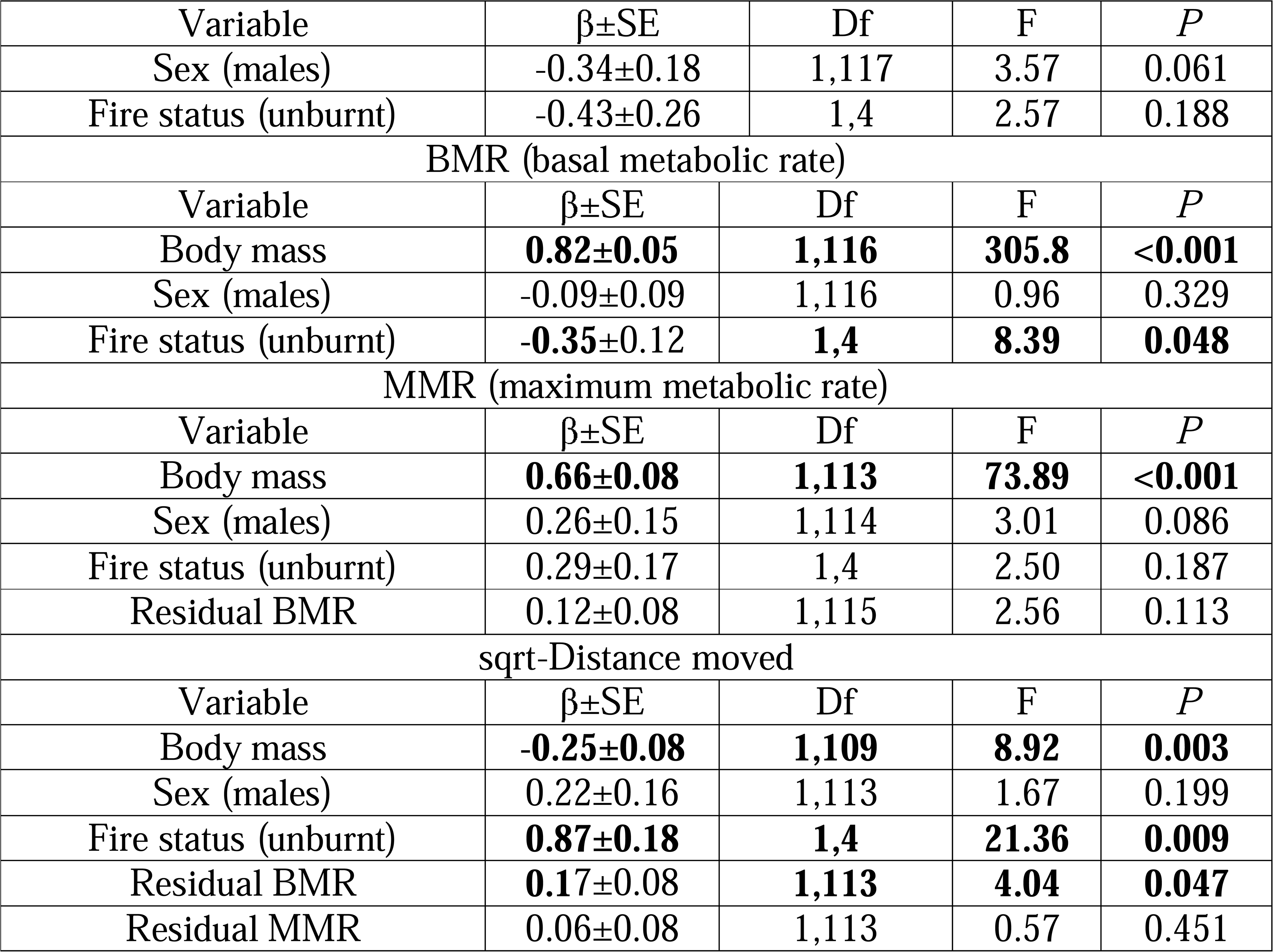
Analysis of variance for body size, basal metabolic rate, maximum metabolic rate and distance moved during open field test. Individual voles were captured at all six study sites located in burnt and unburnt parts of the valley (Fire status).

## Discussion

Only a few studies investigated the effect of fire-related habitat disturbance on animal physiology. Most if not all of them were done in fire-adapted and fire-prone populations of heterothermic animals that are able to enter torpor - a reduction of body temperature that allows for substantial energy savings (review in: Geiser et al., 2018). To the best of our knowledge, no study exists to consider maintenance metabolism in homeothermic animals and its behavioural consequences in fire and other disturbance ecology. Conducted physiological studies concern burrowing or volant species that are able to survive surface fire by hiding below the ground or by flying away. In contrast, we studied fire-naive strictly homeothermic species (Nieminen et al., 2013) with limited movement ability in the habitat (wetlands) where burrowing is not possible.

Our study showed a root vole population collapse in the burnt habitat and a slow recovery a few months after the fire (Fig. 3). We hypothesized that different evolutionary models shape the animal personality in disturbed and undisturbed habitats. We found that physiological traits, maintenance metabolism (BMR) and aerobic capacity for physical exercise (MMR) were repeatable in the studied root vole population (Tab. 1). Distance moved by voles was also repeatable, suggesting that this behaviour (but not the others obtained in this study) is a part of voles personality (Tab. 1). In contrast to what we expected for animals physiology, we did not find an interaction between maintenance (and maximum) metabolism and fire disturbed status of habitat in predicting distance moved by animals (Tab. 2). Instead, we found a consistent positive correlation between maintenance (but not maximum) metabolism and distance moved (Fig. 4a). This result suggests that the ‘performance’ model is dominating personality-physiology association in this species, irrespectively of the presence or absence of the fire disturbance. However, we also recorded a significant increase in the level of BMR, as well as a decrease in the level of mobility of voles, in burnt compared to unburnt habitats (Fig. 4b). The results suggest that besides general physiological performance mechanisms operating in this species, the habitat disturbance might reveal some allocation constraints on physiology-pesionality linkage.

Predation is considered a major driver shaping variation in animal behaviour (Lima and Dill 1990; Dubois and Binning 2022) and the predatory-naive populations may differ in personality traits from those that experience predators (Kortet et al., 2015). Since at post-fire environments, prey species are expected to experience increased predatory pressure (Doherty et al., 2022; Michel et al., 2022) one could argue that observed variation in voles behaviour results from different selection or plasticity in behavioural response to predation (Abbey-Lee et al., 2016). The lower exploration observed in voles captured at post-fire habitat, when compared to those from undisturbed areas, would intuitively support this. Logically, at least at the beginning of recolonization and with reduced grass cover (Fig. 2) the animals in post- fire habitat were probably more exposed to predators and effectively selected for decreased activity than those from unburnt habitat. The post-fire area clearly differed in vegetation cover from the undisturbed part of the habitat during the first month after the fire (Fig. 2). However, the difference in fresh vegetation matter was not detectable on satellite images just two months later (Durka 2020; personal field observations). Thus, even if initially the lack of cover would affect voles behaviour due to increased predatory pressure, it does not necessarily explain observed variation in exploration later in the autumn. Although predation can affect behaviour, the increased maintenance metabolism of voles from post-fire zone when compared to those captured at unburnt habitat cannot be easily explained by predation. The fight- or-flight response requires an elevation of metabolism in response to predators cues (e.g. Lagos and Herberstein 2017), and a comparative study of bird species suggest that high BMR correlate positively with predatory avoidance behaviours (Møller 2009). Yet, the experiments on intraindividual changes in response to predatory cues brings rather weak or no support for this hypothesis (Broggi and Nilsson 2023; see also: Mathot et al., 2016). Moreover, individuals characterized by high self- maintenance metabolism can also be considered more vulnerable to predators due to increased energetic demands, elevated feeding rate and thus a higher probability of becoming prey (Finstad et al., 2007; Krams et al., 2013ab; Mathot et al., 2015). This would result rather in the negative selection on metabolism and related proactive behaviours and the reduction of self-maintenance costs should be expected under high predation.

Body mass (that does not differ between burnt and unburnt habitats) in the studied population of root voles is a good indicator of age and the cohorts overlap in their variation only a little (see: Zub et al., 2014). Distance moved by voles was negatively correlated with among individual variation in *m*_b_, suggesting that younger and more dispersive voles were also more exploratory (Rowell and Rymer 2023).

However, when adjusted for this effect (likely an age-dependent state) the distance moved was a repeatable trait in the studied population (Tab. 1). The size-age independent variation in exploration was positively correlated with consistent among individual variation in BMR (Fig. 3). The high maintenance metabolism reflects developed metabolic machinery needed for the assimilation of energy to improve fitness, consistent with the ‘increased-intake’ hypothesis (Olson 1992; Nelson 2002) and most previous studies on proactivity-BMR association (review in Mathot et al., 2019). However, the ‘aerobic-capacity’ model predicts that individual variation in BMR is a by-product of selection acting on MMR (Hayes and Garland 1995), as those characters are subjected to correlated evolution (Nespolo et al., 2017; but see also: Fiedler and Careau 2021). Since most of the previous studies considered only resting (or basal) metabolism, it remains ambiguous whether variation in behaviour correlate with minimum or rather with maximum metabolism, as suggested by some reviews (Biro et al., 2018). We did not find any association between MMR (that also did not correlate with BMR here) and distance moved in studied voles. This suggests that behavioural expression in root voles (at least in somewhat artificial laboratory settings) is not shaped by aerobic capacity, but rather it is limited by the rate of processing energy to support behavioural performance, in agreement with ‘performance’ models rooted in ‘increase-intake’ hypothesis. Our correlative results on wild animals support results from experiment where artificially selected high BMR resulted in elevated home-cage spontaneous activity in laboratory mice (Gębczyński and Konarzewski 2009a). In the same experiment selection for high MMR during forced swim did not result in any behavioural changes (Gębczyński and Konarzewski 2009b).

There can be several not mutually exclusive reasons for observed elevated BMR in voles in burnt habitat. A ‘food-habits’ hypothesis predicts that environmental productivity shapes the variation in BMR, related to diet variability and quality (review in: Cruz-Neto and Bozinovic, 2004). Note, however, that response to diet quality may differ at ecological and evolutionary time-scales. Since the post-fire environment regrowth vegetation can differ in quality and quantity from the unburnt part of habitat, thus likely corresponding differences may be expected in animal energetics. The vegetation at post-fire areas can be more nutrient-reach (Van de Vijver et al., 1999; Eby et al., 2014) however, at the same time it can also contain more secondary metabolites (Santacruz-Garcia et al., 2021). Both high quality as well as the increased level of secondary compounds in diet can result in elevated metabolism (review in: Cruz-Neto and Bozinovic, 2004). Last but not least, self-maintenance in rodents can strongly respond to thermal conditions (and other stressors), and short to moderate-term acclimation to cold results in reversible elevation of BMR (Boratyński et al., 2016; 2017). Such plastic adjustments of BMR to temperature are rather common among animals and can also be fixed during individual development (reviews in: Swanson et al., 2017; Nornin and Metcalfe 2019). The studied root vole population was suggested to possess high phenotypic plasticity (Zub et al., 2014) and the root voles acclimated to cold are characterized by higher BMR than those acclimated to warm at least when exposed also to short photoperiod (Wang et al., 1999). The root voles built nests in the clumps of sedges composed of dry mater accumulated throughout the years. In result of fire, those shelters were completely burnt (Fig. 2) and the new growth sedges were likely less insulated due to different rations of fresh to dry biomas at burnt and undisturbed habitats (e.g. Eby et al., 2014). Since we measured animals in late autumn when ambient temperature falls below freezing, higher BMR levels in voles from post-fire zone than those from unburnt sites can reflect colder nesting conditions. Nevertheless, if those plausible mechanistic explanations for variation in maintenance metabolism are correct, we found that most likely as compensation voles also varied in behaviour. The higher BMR of voles occupying post-fire zone, when compared to the undisturbed part of habitat, correlated with lower behavioural expression (Fig. 4b) suggesting that when adjusted for positive correlation with metabolism expected by ‘performance’ model (Fig. 4a), the allocation still shaped part of voles behaviour in an additive manner.

## Conclusions

The models based on theory of predatory-prey interactions were offered to explain variation in animal behaviour in post-fire environmet (Doherty et al., 2022; Michel et al., 2022) but they are one side of the coin. Our study suggests that variation in animal personality or behaviour in post-fire habitat can be also shaped by physiological changes as predicted by the intensively studied evolutionary models (Careau et al., 2008; Careau and Garland 2012; Mathot and Dingemanse 2015). In our species the ‘performance’ mechanism seems to have paramount effect on voles behaviour (Fig. 4a). However the ‘compensation’ mechanism is revealed when context of disturbance happen (Tab. 2; Fig. 4b) supporting conclusion that personality-physiology linkage can be situation dependent (Careau and Garland 2012). We suggest that the sometimes-observed lack of a link between behaviour and energetics in numerous previous research (review in: Mathot et al., 2019) could be attributed to both mechanisms masking each other, if the ecological context of the study is not properly addressed (as suggested: Lantová et al., 2011). Both the exploration behaviour and maintenance metabolism were consistent in our study (repeatability of 0.5 and 0.7, respectively), suggesting selection can effectively operate on them. Yet, it cannot be excluded that ‘compensation’ and ‘performance’ mechanisms develop those phenotypes at different, plastic and genetic levels. The correlation between resting metabolism and wheel running can sometimes be positive among-, while negative within-, individual levels (Abdeen et al. 2021). Thus, it can be hypothesised that the ‘performance’ develops phenotype on the genetic, while the ’allocation’, on the plastic levels. Contribution of physiological responses in disturbed (or any stressful) conditions will require experimental design, such as controlled fire, resettlement of individuals between treatments and many repeated measurements. To conclude, post-fire behavioural studies, as well as other behavioural and evolutionary ecology investigations, would benefit from broader application of physiological experiments to tackle changes in energetic environments of animals.

## Acknowledgments

The authors thank Jan R.E. Taylor for allowing us using the laboratories at the Field Station of Faculty of Biology at University of Białystok located in Gugny village for conducting laboratory measurements. The authors also thank Łukasz Ołdakowski for his help during field and laboratory work.

## Data availability

All relevant data can be found within the article and its supplementary information.

